# Single-cell transcriptomics resolves the developmental transitions from schizont to invasive extracellular merozoite to ring-stage *Plasmodium falciparum* parasites

**DOI:** 10.64898/2026.07.27.740906

**Authors:** Sebastiaan Bol, Jake A. Moore, Siddharth Neog, Vinit Upasani, Ian H. Cheeseman, Evelien M. Bunnik

## Abstract

The progression from mature schizonts to newly formed ring-stage parasites represents one of the largest developmental transitions in the *Plasmodium falciparum* intraerythrocytic developmental cycle (IDC), encompassing parasite egress, erythrocyte invasion, and productive infection of the new host cell. While the transcriptional programs required for egress and invasion are well-documented, less is known about how the parasite prepares for intracellular survival inside the newly invaded erythrocyte. Here, we used single-cell RNA sequencing based on PIPseq chemistry to construct a transcriptional atlas spanning late schizogony, extracellular merozoites, and early ring-stage development. We show that schizogony is characterized by sequential waves of invasion gene expression that closely parallel secretory organelle biogenesis and are associated with cooperative regulation by ApiAP2 transcription factors. Immediately prior to egress, parasites shift from expression of invasion genes to genes required for establishing the intracellular host-parasite interface, including the parasitophorous vacuole and digestive vacuole. Extracellular merozoites separate into transcriptionally distinct invasive and non-invasive populations. Together, these findings provide high temporal resolution of late schizont development, reveal priming for intracellular survival prior to egress, and highlight the short lifespan of invasive merozoites and the resulting challenges of studying this elusive stage of the IDC.

## INTRODUCTION

*Plasmodium falciparum* is responsible for the most severe forms of malaria in humans, disproportionately affecting young children in sub-Saharan Africa [1]. Symptoms of malaria are caused by *P. falciparum* parasites in the asexual blood stage, also called the intraerythrocytic developmental cycle (IDC). During each round of the 48-hour IDC, parasites go through three distinct developmental stages. First, during the ring stage, the parasite extensively remodels its host cell by exporting hundreds of proteins into the erythrocyte cytoplasm and synthesizes the translation machinery necessary for the rapid growth phase that follows. Next, during the trophozoite stage, the parasite becomes highly transcriptionally and translationally active to generate the proteins required for cellular replication. Finally, during the schizont stage, the parasite divides into 10–30 daughter merozoites, each equipped with the specialized secretory organelles required for erythrocyte invasion. Fully mature schizonts undergo egress to release merozoites that invade new erythrocytes and continue the replication cycle. The drastic changes in cellular function between these three main developmental stages have been extensively studied and are tightly coordinated by changes in gene expression [2–7]. However, not much is known about how the parasite navigates the transition from a schizont focused on cell division and invasion to a newly formed ring-stage parasite expressing genes involved in erythrocyte remodeling and ribosome biogenesis.

The temporal changes in gene expression during the schizont-to-ring transition have been difficult to characterize for two reasons. First, the synchronization window that can be achieved by standard parasite culture methods is too broad to accurately separate fully segmented schizonts from early ring-stage parasites. Bulk RNA sequencing (RNA-seq) approaches are therefore unable to resolve the transition in transcriptional programs between these developmental stages. Second, substantial transcriptional remodeling in the hours before egress has led to under-sampling of late-stage schizonts and merozoites in single-cell RNA-seq experiments [8].

To partially overcome these issues, we previously isolated merozoites by treating early schizonts with E64, a protease inhibitor that prevents rupture of the erythrocyte membrane and thus egress of matured merozoites [9,10]. E64-treated schizonts progress through segmentation normally. Forcing the fully segmented schizonts through a 1.2-µm filter results in mechanical rupture of the schizont membrane, releasing individual merozoites [11]. Bulk RNA-seq of merozoites isolated following E64 treatment revealed substantial transcriptomic heterogeneity between biological replicates [12]. This observed heterogeneity could reflect true developmental variation and be the result of variations in timing, suggesting that merozoites are transcriptionally active with highly dynamic gene expression profiles. Alternatively, it could be a technical artifact arising from differences in the conditions during sample collection. Merozoites are short-lived and their half-life is temperature dependent [11]. Even minor differences in sample handling may therefore introduce transcriptomic variation between samples. These technical challenges have hindered efforts to comprehensively characterize the transcriptomic landscape during *P. falciparum* egress and invasion.

In this study, we aimed to resolve the transcriptomic changes that drive the schizont-to-ring transition in *P. falciparum* blood-stage parasites. To do so, we used a novel single-cell RNA-seq platform called Pre-templated (or Particle-templated) Instant Partition sequencing (PIPseq) [13]. PIPseq allows the encapsulation of individual cells in uniform droplet emulsions using only a vortexer. This approach captures all cells simultaneously and minimizes the time between sample collection and cell partitioning, in contrast to microfluidics systems in which cells are captured sequentially. We hypothesized that this method would preserve the transcriptomic state of cells that undergo rapid changes in gene expression and thus allow for capture of highly transient stages of parasite development during the late schizont, merozoite, and early ring stages. We organized our analysis into three complementary areas. First, we established a temporal map of the *P. falciparum* IDC and characterized changes in transcriptome complexity throughout parasite development. Second, we defined the timing and transcription factor regulation of invasion gene expression during schizont maturation. Third, we examined the transcriptional identity of extracellular merozoites and the effects of pharmacological inhibition of egress through comparison of naturally egressed and E64-treated parasites. Our analyses provide a high-resolution atlas of parasite development from late schizonts to early ring-stage parasites and yield new biological insights into the transcriptional programs that govern this highly dynamic phase of the IDC.

## RESULTS

### PIPseq enables single-cell transcriptomic profiling of *P. falciparum* parasites

To provide proof-of-concept for use of the PIPseq platform to analyze single-cell transcriptomes of *P. falciparum* parasites during the IDC, we generated a single library of an unsynchronized *P. falciparum* 3D7 blood-stage culture. The PIPseq platform successfully captured the transcriptomes of parasites in all stages of the IDC, as well as early and late gametocytes (**Fig. S1A**). To compare these PIPseq results with data obtained using the 10x Genomics platform, we integrated our PIPseq library with a single-cell RNA-seq data set of mixed-stage *P. falciparum* 3D7 parasites from the Malaria Cell Atlas [14] (**Fig. S2**). The integrated data set was visualized by Uniform Manifold Approximation and Projection (UMAP), in which cells with similar transcriptional profiles are positioned together in two-dimensional space (**Fig. 1A**). After removing cells corresponding to gametocytes (**Fig. S1B**) and a small cluster with cells of unknown stages, which are possible doublets, we obtained transcriptomes for 2,712 and 6,267 asexual blood-stage parasites in the PIPseq and 10x Genomics data sets, respectively with medians of 416 and 344 genes detected per cell (**Fig. S3**). The combined UMAP recreates the distinct cyclical nature of the IDC and shows high concordance between the two data sets (**Fig. S1C**). These results demonstrate that PIPseq is a suitable single-cell RNA-seq platform for *P. falciparum*.

**Figure 1:**
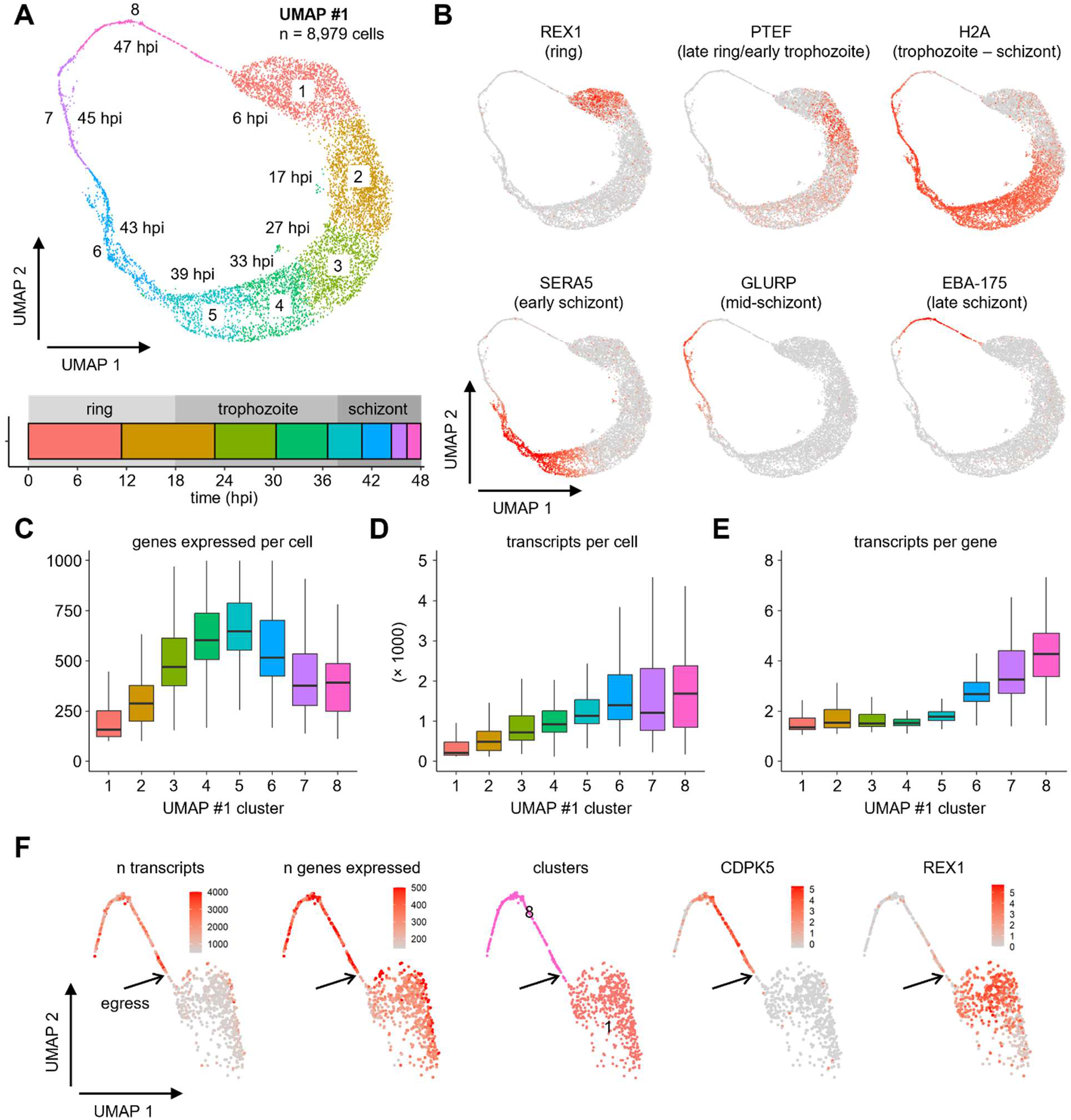
Single-cell RNA-seq analysis of the *P. falciparum* intraerythrocytic developmental cycle enables temporal mapping and identification of the moment of egress. **A)** Integrated UMAP #1 of two scRNA-seq data sets of mixed-stage *P. falciparum* strain 3D7 cultures, prepared using the 10x Genomics and PIP-seq platforms. The graph in the bottom indicates the estimated timing in hours post-invasion (hpi) for each cluster. **B)** Expression of select stage-specific genes projected onto UMAP #1: ring-exported protein 1 (REX1), translation-enhancing factor (PTEF), histone H2A, serine repeat antigen 5 (SERA5), glutamate-rich protein (GLURP), and erythrocyte binding antigen-175 (EBA-175). **C)** The number of genes expressed in each cell per UMAP #1 cluster. **D)** The number of transcripts detected in each cell per UMAP #1 cluster. **E)** The number of transcripts detected per expressed gene in each cell per UMAP #1 cluster. **F)** The number of transcripts, the number of expressed genes, and expression of the late schizont gene CDPK5 and early ring-stage gene REX1 for each cell projected onto clusters 1 and 8 of UMAP #1 containing late schizonts to early rings. The arrow indicates the moment of egress.

### Single-cell transcriptomes enable temporal mapping of the intraerythrocytic developmental cycle

To facilitate biological interpretation of transcriptional profiles, we inferred the developmental timing of each cluster across the IDC. We assumed that the PIP-seq and 10x Genomics data sets were generated from fully asynchronous parasite cultures, such that parasites from all time points throughout the IDC were equally represented. Under this assumption, a large cluster represents a longer time frame than a small cluster. Based on the number of cells per cluster, we calculated the estimated midpoint of each cluster in hours post-invasion (hpi) (**Fig. 1A**, **Table S1**). This showed that ring (0 – 18 hpi) and trophozoite (18 – 38 hpi) stage parasites together occupied the first half of the UMAP circle and were divided into four clusters (clusters 1 – 4, **Fig. 1A**). Schizonts (38 – 48 hpi) were spread out over the remaining four clusters containing decreasing numbers of cells (clusters 5 – 8, **Fig. 1A**). These stage annotations closely matched those of the Malaria Cell Atlas (**Fig. S1D**). As an additional means to validate these timing estimates, we plotted the expression of representative highly stage-specific genes onto the UMAP (**Fig. 1B**). The first cluster was enriched for exported proteins, represented by ring-exported protein 1 (REX1), in line with a focus on erythrocyte remodeling in ring-stage parasites. In the second cluster, genes related to translation were upregulated, as observed in late rings and early trophozoites, represented by translation-enhancing factor (PTEF). Next, genes encoding histones were upregulated in mid-trophozoites in preparation for genome replication and were expressed throughout schizogony (histone H2A). Finally, schizonts sequentially expressed genes encoding proteins involved in erythrocyte egress and invasion, including serine repeat antigen 5 (SERA5), glutamate-rich protein (GLURP), and erythrocyte binding antigen 175 (EBA-175). Together, these findings demonstrate that our analysis provides a reliable temporal annotation of single-cell transcriptomes across the IDC.

### Transcriptome complexity changes throughout the intraerythrocytic developmental cycle

The unequal distribution of parasite stages along the developmental trajectory suggests that the extent of transcriptomic remodeling varies across the IDC. To investigate the basis of this observation, we quantified the number of expressed genes and transcripts per cell in each cluster. The number of expressed genes per cell gradually increased during ring and trophozoite development, peaked in early schizonts (cluster 5), and then gradually decreased again (**Fig. 1C**). The number of transcripts per cell, on the other hand, continued to increase throughout the IDC until it plateaued in schizonts (clusters 6 – 8) (**Fig. 1D**). Because sequencing depth is an important determinant of both the number of transcripts captured and number of genes detected, we explored the relationship between both metrics for each cluster. Although sequencing depth explained most variation in detected genes, clusters showed differences in the relationship between the two metrics even after adjusting for sequencing depth, indicating that cell clusters differed in transcriptome complexity (**Fig. S4**). The average number of transcripts per expressed gene was relatively stable during the ring and trophozoite stages, but increased in schizonts until it peaked at the end of the IDC (**Fig. 1E**). This increase in transcripts per gene was independently observed in both data sets (**Fig. S5**) and may reflect multiplication of the *P. falciparum* genome during schizogony. During this process, transcription from multiple gene copies may be necessary to meet the demands of cell division and merozoite formation. Collectively, these data reveal that transcriptome complexity changes throughout the IDC, with schizont maturation characterized by increased transcript output from a more restricted set of expressed genes.

### Single-cell RNA-seq analysis enables identification of the moment of egress

One of the primary goals of our study was to characterize the transcriptional changes that occur during the final moments of the IDC, as segmented schizonts rupture, releasing merozoites that invade new erythrocytes and initiate ring-stage development. A major challenge in addressing this question is the accurate identification of the egress event. We hypothesized that the transition from multinucleated segmented schizonts to individual merozoites would be associated with a sharp decline in the number of transcripts and genes detected per cell, providing a transcriptomic signature of egress. To analyze this, we extracted the late schizont stage and early ring stage clusters (1 and 8 in **Fig. 1A**) from UMAP #1 and projected the two relevant metrics (transcript count and gene count) for each cell onto the resulting UMAP. An abrupt decrease in both transcript content and the number of expressed genes detected per cell was visible close to the border between the two clusters (**Fig. 1F**), suggesting this is the moment that merozoites are released from the erythrocyte. This designation also fit with changes in gene expression during the transition from schizont to ring. Expression of the gene encoding calcium-dependent protein kinase 5 (CDPK5), which is essential for the release of merozoites from the host cell [15], was dramatically reduced upon egress. This was immediately followed by upregulation of the early ring-stage gene REX1. Together, these results indicate that the transition from fully segmented schizonts to newly formed rings is marked by a sharp contraction of the transcriptional landscape and a shift in gene expression program, consistent with parasite egress and the onset of a new IDC. Of note, the transition between schizont and ring-stage parasites appears to be incorrectly annotated in several *P. falciparum* data sets available from the Malaria Cell Atlas (**Fig. S6**) and the associated publication [16]. This observation underscores the importance of accurately determining the moment of egress to interpret stage-specific gene expression and distinguish pre-egress from post-invasion transcriptional programs.

### A schizont-enriched PIPseq data set increases resolution of late IDC development

Although PIPseq captured all stages of the IDC, cells in the mid-to-late schizont stages and extracellular merozoites were sparsely sampled. To resolve changes in gene expression during the schizont-to-ring transition at higher resolution, we generated an additional PIPseq library from a culture enriched for schizonts, merozoites, and early ring-stage parasites (see Methods for details). This library was integrated with the PIPseq and 10x Genomics libraries of mixed-stage *P. falciparum* cultures to enable temporal annotation of cells throughout the IDC and visualized in UMAP #2 (**Fig. S2, S7A**). After removing a cell cluster containing gametocytes and four small clusters that were not positioned along the circular trajectory of UMAP #2, the combined data set contained transcriptomes of 12,882 parasites divided over 11 clusters (**Fig. 2A**). Using cells from the PIPseq and 10x Genomics mixed-stage *P. falciparum* libraries to estimate the timing in hours post-invasion for each cluster, we determined that clusters 1 and 2 contained rings (0 – 18 hpi) and clusters 3 and 4 contained trophozoites (18 – 38 hpi). Cluster 5 was a mix of late trophozoites and early schizonts, while the remaining six clusters were made up of schizonts and merozoites (40 – 48 hpi) (**Fig. 2A**, **Table S1**). The large majority of cells (88%) in the library enriched for schizonts, merozoites, and rings were present in clusters 1 (early rings) and 6 – 11 (schizonts and merozoites) as opposed to <40% in the two mixed-stage samples (**Fig. S7B**, **Table S1**). Moreover, UMAP #1 cluster 8 harbored 313 cells, which covered approximately the last two hours of schizogony. The same time frame is represented by 1,716 cells divided over three clusters in UMAP #2 (**Fig. S7C**, **Table S1**). These results confirm that our enrichment strategy was successful and that we had achieved higher resolution of clusters covering the late schizont stage.

**Figure 2:**
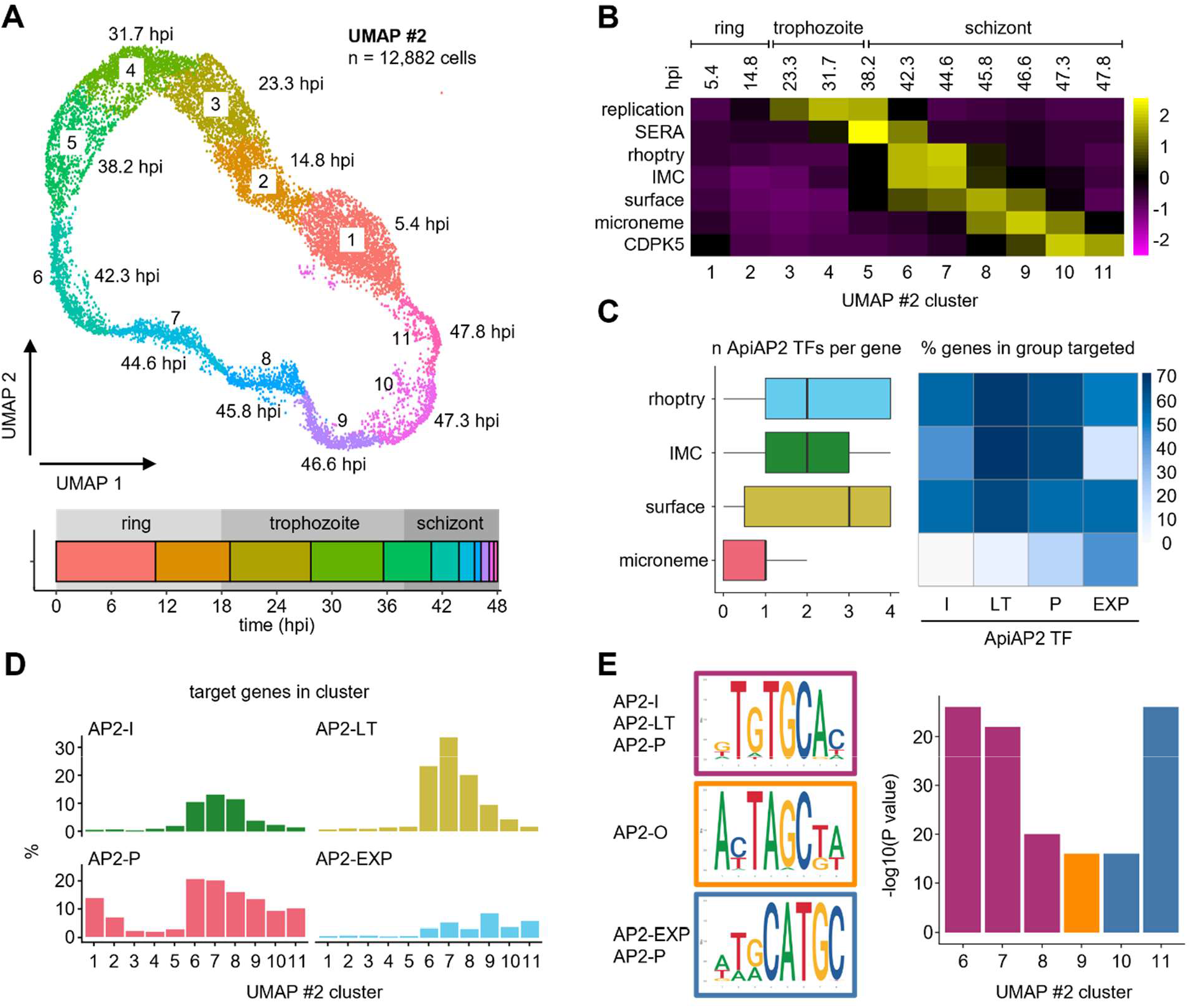
Invasion genes are expressed in sequential waves during schizogony. **A)** Integrated UMAP #2 of three scRNA-seq data sets: two mixed-stage *P. falciparum* strain 3D7 cultures and a culture enriched for schizonts, merozoites, and early rings. The graph in the bottom indicates the estimated timing in hours post-invasion (hpi) for each cluster. **B)** Average expression of modules of genes involved in replication and merozoite formation during the IDC. **C)** The number of ApiAP2 transcription factors (TFs) targeting different group of invasion genes (left) and the percentage of genes in each group of invasion genes that is bound by AP2-I, AP2-LT, AP2-P, and AP2-EXP (right). **D)** The percentage of genes in each UMAP #2 cluster bound by AP2-EXP, AP2-I, AP2-LT, and AP2-P. **E)** DNA binding motifs enriched in promoters of genes with peak expression in the schizont clusters of UMAP #2 (6 – 11). SERA, serine repeat antigen; IMC, inner membrane complex; CDPK5, calcium-dependent protein kinase 5.

### Invasion genes are expressed in sequential waves during schizogony

We first focused our analysis on transcriptional changes during schizont maturation. Poran *et al*. [17] previously reported that gene sets encoding proteins involved in merozoite egress and invasion are expressed in waves during schizogony. In that analysis, gene expression was plotted along an axis of arbitrary pseudo-time units. To estimate the actual timing of gene expression, we calculated relative expression of the gene modules from Poran *et al.* across all clusters and added a new gene module containing merozoite surface proteins (**Tables S2, S3**). This analysis recapitulates the sequential expression of these gene sets during schizogony, starting with genes encoding serine repeat antigen (SERA) proteins in early schizonts (38 hpi), followed by genes encoding rhoptry and inner membrane complex proteins in mid-schizonts (42 – 45 hpi), merozoite surface proteins in mid to late schizonts (44 – 47 hpi), microneme proteins in late schizonts (46 – 47 hpi), and concluding with the expression of CDPK5 in the final hour prior to egress (47 – 48 hpi) (**Fig. 2B, Fig. S8**). These results confirm the sequential expression of invasion-related gene modules during schizogony and define the timing of the transcriptional cascade that accompanies merozoite maturation.

### ApiAP2 regulatory control shifts in late schizogony

Gene expression in *P. falciparum* is in part controlled by the family of 27 Apicomplexan Apetala AP2 (ApiAP2) transcription factors [18–20]. To determine whether ApiAP2 transcription factors are involved in regulating the transcriptional cascade of invasion genes, we assessed which ApiAP2 transcription factors are associated with each invasion-related gene module. Several ApiAP2 transcription factors have been implicated in regulating invasion gene expression, including AP2-I (invasion), AP2-LT (late trophozoite), AP2-P (pathogenesis), and AP2-EXP (export). Using published lists of genes bound by each of the four ApiAP2 transcription factors defined by chromatin immunoprecipitation sequencing (ChIP-seq) during the intraerythrocytic schizont stage [21–24], we observed substantial overlap in their target genes, with multiple ApiAP2 transcription factors regulating the same invasion genes. Except for SERA5, the egress-related SERA family genes were not bound by these ApiAP2 transcription factors (**Table S2**). In contrast, most genes encoding invasion proteins that localize to the rhoptries, inner membrane complex, or surface were shared targets of two or more ApiAP2 transcription factors, with AP2-I, AP2-LT, and AP2-P contributing comparably to their regulation (**Fig. 2C**). Genes encoding rhoptry and merozoite surface proteins were also bound by AP2-EXP. Genes encoding microneme proteins, expressed later than the other three groups of genes, showed a distinct pattern. Of the four ApiAP2 transcription factors, only AP2-EXP bound a substantial fraction of microneme genes (43%; **Fig. 2C**). Together, these findings support a model in which cooperative ApiAP2 transcription factor binding fine-tunes gene expression during schizogony, generating distinct temporal waves corresponding to proteins with different subcellular localizations.

The observation that microneme genes are expressed later than other invasion genes and were primarily bound by AP2-EXP prompted a broader analysis of ApiAP2 transcription factor binding patterns during schizogony. For each UMAP #2 cluster, we quantified the fraction of genes with peak transcript abundance that are bound by each of the four ApiAP2 transcription factors. Consistent with peak expression of invasion-associated genes in cluster 6 – 9, approximately one third (23 – 38%) of genes in these cluster were bound by AP2-I, AP2-LT, AP2-P, or AP2-EXP, with the most prominent contributions by AP2-LT and AP2-P, followed by AP2-I (**Fig. 2D**). In the final two clusters of the IDC (clusters 10 and 11), the proportion of AP2-I and AP2-LT target genes substantially decreased, such that the largest fraction of genes was now bound by AP2-P or AP2-EXP (**Fig. 2D**), indicating a shift in transcriptional control as parasites reach full maturation.

To validate this regulatory transition, we assessed whether promoter regions of genes peaking in each cluster were enriched for DNA binding motifs of ApiAP2 transcription factors. In agreement with the analysis of target genes identified by ChIP-seq, clusters 6 – 8 were enriched for the GTGCA motif, recognized by AP2-I, AP2-LT, and AP2-P (**Fig. 2E**). In contrast, cluster 9 showed enrichment for the TAGCTA motif associated with AP2-O, which is known to regulate microneme genes in *P. berghei* ookinetes [25,26] and may have a similar role in the *P falciparum* IDC. Finally, clusters 10 and 11 showed enrichment for the GCATGC motif, a secondary AP2-P motif shared with AP2-EXP (**Fig. 2E**). Together, these findings suggest that while AP2-P acts in concert with AP2-I and AP2-LT during earlier stages of schizogony, regulatory control shifts to AP2-O and finally to AP2-P and AP2-EXP during the final phases of the IDC.

### Extracellular merozoites separate into transcriptionally distinct invasive and non-invasive populations

Because E64 is commonly used to isolate merozoites, we next evaluated whether E64 treatment altered the transcriptional landscape during the schizont-to-ring transition. We generated two additional PIPseq libraries of E64-treated parasites, one containing schizonts sampled directly from the culture and one containing merozoites obtained following mechanical rupture of E64-treated schizonts. We then combined data from all five samples (PIPseq and 10x Genomics libraries of mixed-stage *P. falciparum* parasites, the library enriched for schizonts, merozoites, and rings, and the libraries of E64-treated schizonts and merozoites; **Fig. S2**). The integrated UMAP again showed the familiar circular pattern with a separate cluster of what appeared to be individual merozoites (**Fig. S9**). The five clusters representing late schizonts, merozoites, and early ring-stage parasites were extracted and re-clustered to obtain higher resolution of the developmental trajectory surrounding egress and invasion (UMAP #3, **Fig. 3A**).

**Figure 3:**
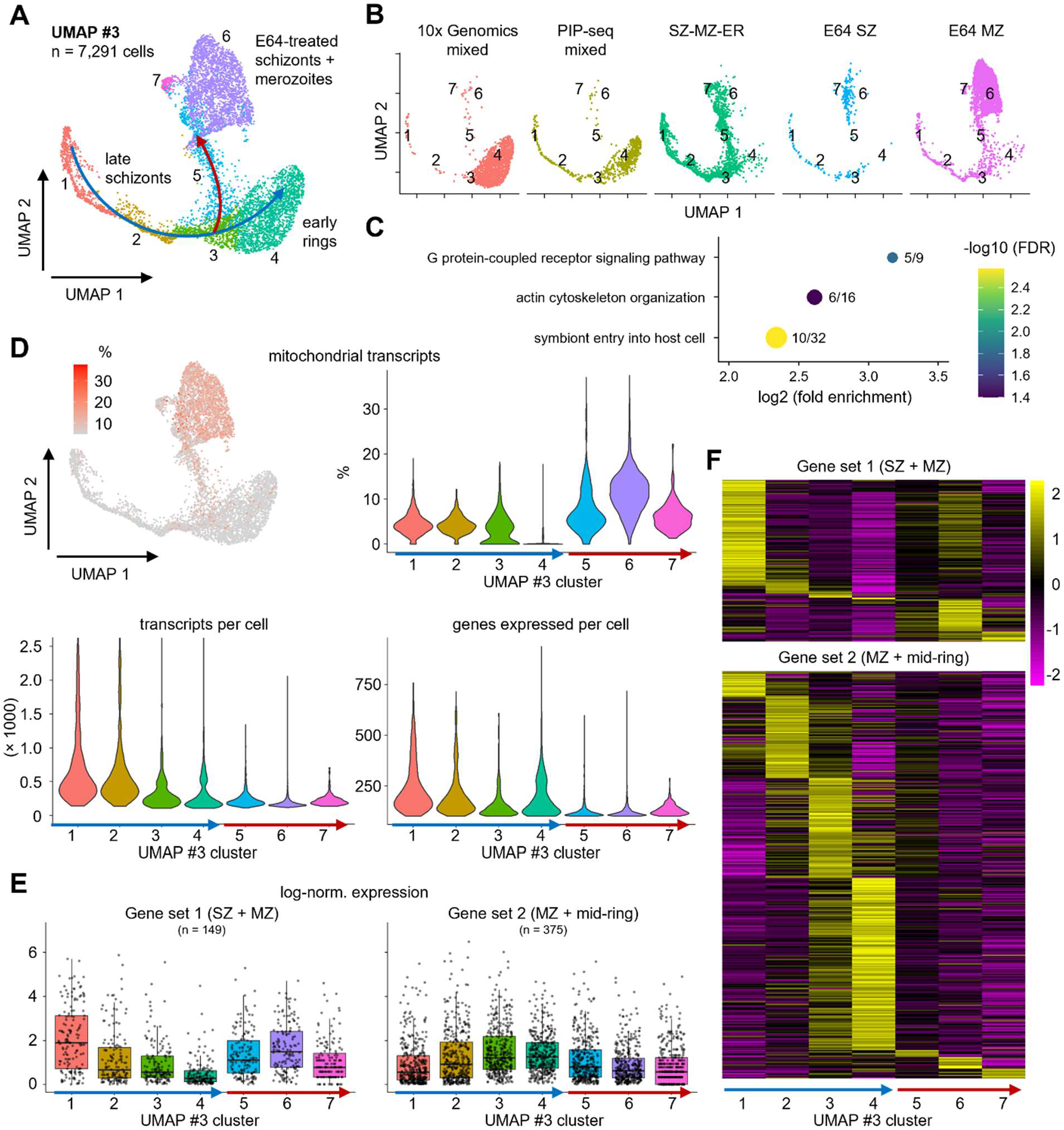
Extracellular merozoites separate into transcriptionally distinct invasive and non-invasive populations. **A)** Integrated UMAP #3 containing cells of five scRNA-seq data sets: two mixed *P. falciparum* strain 3D7 cultures, a culture enriched for schizonts, merozoites, and early rings (SZ-MZ-ER), E64-treated schizonts (E64 SZ), and merozoites obtained by mechanical rupture of E64-treated schizonts (E64 MZ). The normal developmental trajectory and diverging branch are indicated with blue and red arrows, respectively. **B)** The same UMAP #3 shown in panel A split by sample of origin. **C)** Gene ontology pathways enriched among genes with elevated expression in the diverging branch (clusters 5 – 7) relative to the normal developmental trajectory (clusters 1 – 4). Circles are color-coded according to false discovery rate (FDR) and sized based on the number of genes associated with the gene ontology term. The fraction of genes found for each gene ontology term is indicated. **D)** The percentage of mitochondrial transcripts projected onto UMAP #3 (top left) and plotted for each cluster (top right). The bottom row shows the number of transcripts detected in each cell (left) and the number of genes expressed in each cell (right) per UMAP #3 cluster. **E)** Expression of genes in gene set 1 and 2 from ref. [12] in UMAP #3 clusters. **F)** Relative expression of gene sets 1 and 2 [12] in UMAP #3 clusters.

Data from the two mixed-stage cultures were used to delineate the normal developmental pathway spanning late schizont to early ring-stages. In the 10x Genomics mixed-stage sample, the late schizont trajectory (clusters 1 and 2) was not fully connected and the majority of parasites resided in the early ring-stage cluster 4 (**Fig. 3B**, left, **Table S1**). In contrast, the mixed-stage PIPseq sample exhibited a more balanced stage distribution and a continuous late schizont-stage trajectory without observable gaps (**Fig. 3B**, center-left, **Table S1**). In both samples, a minor subset of cells did not align with the principal developmental trajectory, and were positioned along a distinct branch consisting of clusters 5 – 7.

As expected, the schizont-merozoite-ring library was enriched for late schizont-stage parasites (clusters 1 and 2; **Fig. 3B**, center). However, only a minority of parasites from this sample mapped onto early ring-stage cluster 4 in the primary developmental trajectory defined by the mixed-stage samples. Instead, the majority of parasites populated the diverging branch consisting of clusters 5 – 7. While this branching did not occur in UMAP #2 that covered the entire IDC, a similar organization was visible when the late schizont and early ring clusters were extracted and re-clustered (**Fig. S10**). Cells from both E64-treated samples were also predominantly located in the diverging branch (**Fig. 3B**, center-right and right). Together, these observations suggest that both naturally and mechanically released merozoites initiate progression along this diverging trajectory, most likely due to prolonged residence in the extracellular environment or delayed egress. Cells along this divergent branch ultimately accumulate at a distinct terminal state that may represent merozoites that are no longer invasive. Notably, the mechanically released E64-treated merozoite sample also contained late-stage schizonts (∼10% of cells, **Table S1**) despite passage through a 1.2-µm filter and subsequent negative magnetic selection. This finding underscores the inadequacy of these methods to completely eliminate residual schizonts and the need for caution when interpreting data derived from mechanically isolated merozoite preparations.

To better understand the transcriptional identity of merozoites in the diverging branch, we analyzed genes that were differentially expressed between clusters 5 – 7 and the normal developmental trajectory (cluster 1 – 4). A total of 1,542 genes differed in expression by more than two-fold, with only 342 genes showing higher expression in cells of the diverging branch (**Table S4**). Gene ontology analysis of these 342 genes revealed five significantly enriched biological processes (false discovery rate < 0.05), including G protein-coupled receptor signaling, actin cytoskeleton organization, and symbiont entry into host (**Fig. 3C, Table S5**). Representative genes associated with these gene ontology annotations included cAMP-dependent protein kinase catalytic subunit (PKAc), myosin E (myoE), and merozoite surface protein 2 (MSP2) (**Fig. S11**). In addition to these nuclear genes, 34 mitochondrial genes were more highly expressed in parasites in the divergent branch, whereas none were downregulated. An increased proportion of mitochondrial transcripts is a sign of cell damage resulting from loss of cytoplasmic mRNA. Indeed, cells along the divergent branch showed increased percentages of mitochondrial transcripts, lower overall transcript abundance, and fewer detected genes (**Fig. 3D**), consistent with cellular damage. Taken together, these findings suggest that the transcriptional profile of non-invasive merozoites reflects cellular deterioration with a remnant of the late schizont-stage gene expression program.

In a previous bulk RNA-seq study, we identified 655 genes that were upregulated in merozoites obtained by E64 treatment and subsequent mechanical rupture, relative to early schizonts collected at 40 hpi [12]. To place these findings in the context of our new observations suggesting that the majority of E64-treated merozoites have a distinct transcriptional profile from invasive merozoites, we examined the expression of these genes along the developmental trajectory. These genes had previously been separated into two distinct groups based on their expression profile. The first group consisted of genes (n = 205) upregulated in late schizonts and merozoites. Most of these genes peaked in UMAP #3 cluster 1, corresponding to late schizonts (62%), with progressively decreasing expression in subsequent late-schizont and early-ring clusters (clusters 2 – 4) (**Fig. 3E, F, Table S6**). Expression of this first gene set was also elevated in merozoites along the diverging branch, with 20% of genes peaking in cluster 6. These results suggest that this gene set primarily captured both contaminating schizonts and non-invasive merozoites. The second group consisted of genes (n = 450) with high expression in merozoites and mid-rings, with substantial variation between biological replicates of merozoite samples [12]. In UMAP #3, most of these genes peaked in clusters 3 and 4, which encompass free merozoites and early rings, consistent with transcriptional programs associated with normal egress and re-invasion (**Fig. 3E, F, Table S6**). This second gene set thus most accurately captured genes upregulated during merozoite formation and early ring-stage development. The observation that merozoite samples in our prior analysis [12] showed highly variable expression of these genes points towards substantial batch-to-batch variation in merozoite quality.

### Late schizonts and merozoites deploy a staged transcriptional program to establish the intracellular host-parasite interface

The normal developmental trajectory in UMAP #3 spans the last two hours of schizogony and the first four hours of ring-stage development. To analyze changes in gene expression during this time span in more detail, we first identified the moment of egress in UMAP #3 based on the sharp decline in expression level of late-schizont gene CDPK5 (**Fig. 4A**), as established in UMAP #1. We then clustered the cells at a higher resolution to obtain seven clusters along the normal developmental trajectory (UMAP #3 High Resolution (HR); **Fig. 4B**). Cluster 4 consisted almost exclusively of free merozoites, defined as cells that no longer expressed CDPK5 and did not yet express the early-ring gene REX1 (**Fig. 4A**).

**Figure 4:**
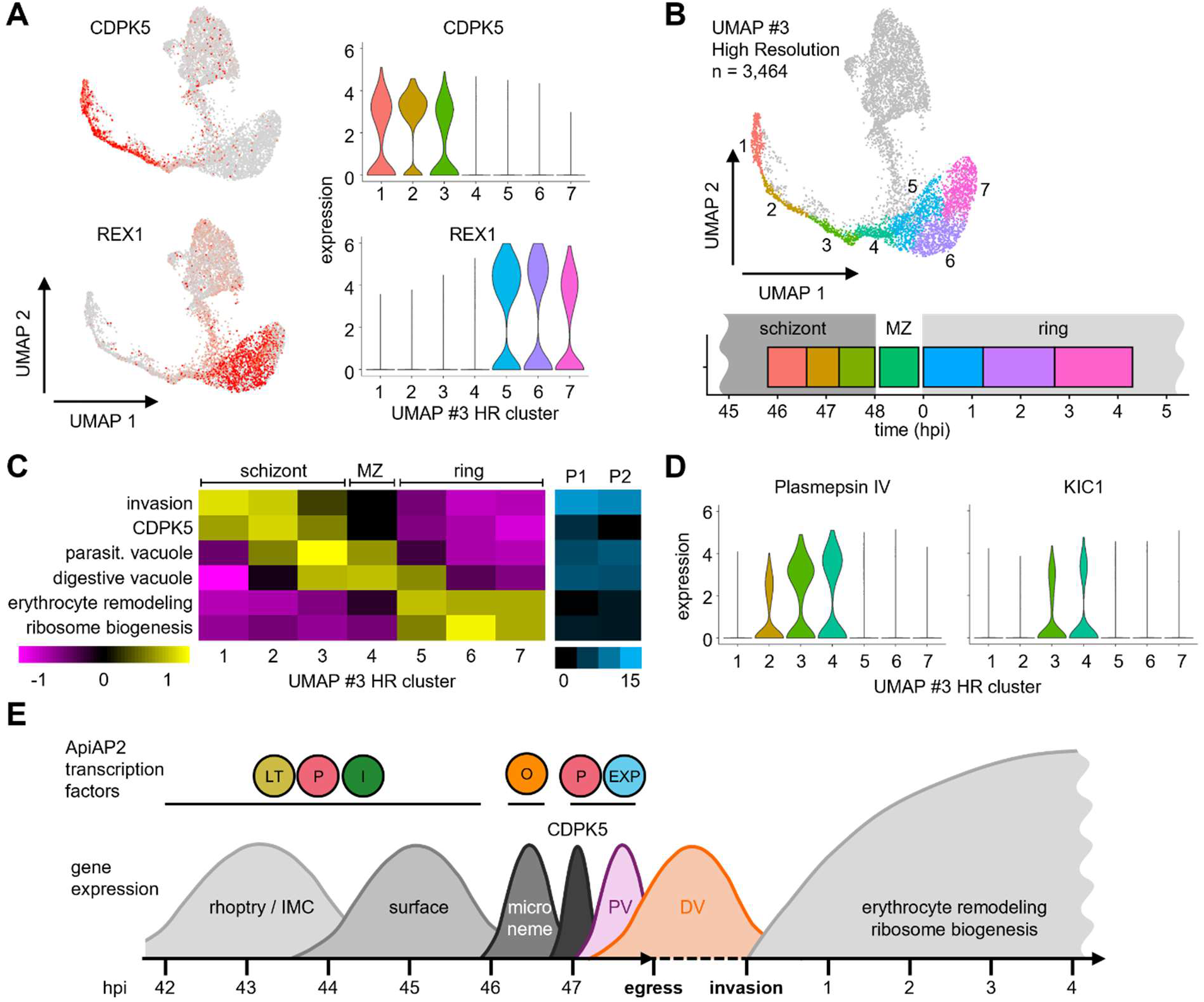
Just-in-time gene expression during the schizont to ring transition. **A)** Expression of late-schizont gene CDPK5 and early ring-stage gene REX1 projected onto cells of UMAP #3 (left) and as violin plot showing clusters along the normal developmental trajectory in high-resolution UMAP #3 (right). **B)** UMAP #3 clustered at a higher resolution (0.7). Cells along the normal developmental pathway are colored (clusters 1 - 7), while cells in the diverging branch are shown in gray. The graph in the bottom indicates the estimated timing in hours post-invasion (hpi) for each cluster. **C)** Average relative expression of modules of genes involved in merozoite formation and early-ring stage development during the schizont-to-ring transition. On the left, the average number of unique peptides per gene is depicted for two independent proteomics studies of merozoites. **D)** Two example genes that peak in expression in merozoites (UMAP #3 HR cluster 4): plasmepsin IV and Kelch13-interacting candidate 1 (KIC1). **E)** Model of gene expression during the schizont-to-ring transition. Changes in ApiAP2 transcription factor targeting contribute to waves of merozoite gene expression during the final hours of the intraerythrocytic developmental cycle of *P. falciparum*, culminating in the expression of core components of the parasite-erythrocyte interface: the parasitophorous vacuole (PV) and digestive vacuole (DV). Following erythrocyte invasion, the transcriptional landscape shifts to the expression of genes involved in, among other processes, erythrocyte remodeling and ribosome biogenesis. MZ, merozoite.

For each of the seven cluster along the late schizont to early ring-stage trajectory, we identified a list of genes that were both upregulated in differential gene expression analysis and peaked in expression in that particular cluster (**Table S4**). Cells in cluster 1 expressed genes involved in invasion, including EBA-140 and EBA-175, and reticulocyte binding protein homologue 5 (RH5) (**Fig. 4C, S12, S13**). In cluster 2, CDPK5 peaks in expression, just in time for its essential function in merozoite egress. In the very last moments before egress (cluster 3), the parasite expresses components of the *Plasmodium* translocon of exported proteins (PTEX) complex that transports proteins from the parasitophorous vacuole to the erythrocyte cytoplasm (PTEX88, PTEX150, heat shock protein 101 [HSP101], and parasitophorous vacuolar proteins 1 [PV1] and 3 [PV3]). Genes encoding additional proteins located in the parasitophorous vacuole also reach peak transcript abundance, including P113, liver stage antigen 3 [LSA3], and early transcribed membrane proteins (ETRAMPs) 2, 11.1, 11.2, and 12 (**Fig. S12**). These results suggest that just before parasite egress, *P. falciparum* prepares for infection of a new erythrocyte by shifting its transcriptional program from merozoite formation to core components of the parasitophorous vacuole. This parasite structure forms the main interface between the parasite and the host cell and is essential for establishing a suitable environment for the parasite to survive within the host cell.

The subsequent merozoite cluster 4 showed peak abundance of transcripts encoding proteins involved in hemoglobin uptake and digestion, including plasmepsin IV, Kelch13-interacting candidates (KIC) 1 and 7, and multidrug resistance protein 1 (MDR1) (**Fig. 4D**). While the timing of digestive vacuole gene expression was more variable than that of parasitophorous vacuole genes (**Fig. S13**), it converged on a maximum in merozoites. The majority of proteins encoded by genes in the parasitophorous vacuole and digestive vacuole gene sets were detected in the merozoite proteome (**Fig. 4C**, **Table S3**). Beyond the digestive vacuole, merozoites also showed peak expression of gene modules implicated in transcriptional regulation (bromodomain protein 2 and nucleosome assembly protein) and post-transcriptional regulation (polyadenylate-binding protein 1 [PABP1], polyadenylate-binding protein-interacting protein 1 [PAIP1], DNA/RNA-binding protein Alba 4, and RNA-binding protein Musashi) (**Fig. S12**). More than 90% of cluster 4 genes were also upregulated in the preceding late schizont cluster, suggesting that merozoites have minimal *de novo* transcriptional activity.

Early ring-stage parasites (clusters 5 – 7) exhibited a distinct transcriptional program focused on erythrocyte remodeling and ribosome biogenesis (**Fig. 4C**), with expression of various exported proteins and genes encoding ribosome proteins (**Table S4**). Many of these genes were upregulated in cluster 5, even when their abundance peaked in cluster 6 or 7 (**Fig. S13**). Collectively, these results suggest that as *P. falciparum* winds down gene expression just prior to egress, the parasite briefly turns on genes that shape the intracellular host-parasite interface, including the parasitophorous vacuole and the digestive vacuole (**Fig. 4E**). This is followed by a switch to a larger and more sustained gene expression program shortly after invasion.

## DISCUSSION

Merozoites are an elusive stage of the *Plasmodium* intraerythrocytic developmental cycle due to their short-lived nature and the technical challenges of isolating pure populations of viable cells. Here, we used single-cell RNA sequencing to characterize the transcriptional landscape during schizont development and merozoite formation. We find that the transcriptomic landscape during schizogony is defined by sequential waves of invasion gene expression, likely coordinated by cooperative ApiAP2 transcription factor activity. Following this invasion gene cascade, genes essential for establishing the intracellular host-parasite interface are upregulated just prior to merozoite egress. Together, these transcriptional changes underlie the successful transition of mature parasites into a newly invaded host cell and prime the parasite for productive erythrocyte infection.

The findings presented here refine and extend previous studies of gene expression throughout the *P. falciparum* IDC [2–5] by providing a higher-resolution view of the transcriptional events leading up to merozoite egress. Although several additional high-resolution bulk RNA-seq data sets are now available on PlasmoDB [27,28], incomplete parasite synchronization and subtle differences in developmental staging between studies have limited the precise assignment of gene expression timing. scRNA-seq overcomes these hurdles by enabling computational ordering of individual parasites along a continuous developmental trajectory. This approach allowed us to estimate the timing of peak transcript abundance at approximately 2-hour resolution during the early-to-mid schizont stage, and at sub-hour resolution in late schizonts, where transcriptional changes are most rapid.

It is important to note that our analysis is primarily based on the timing of peak transcript abundance rather than transcriptional activity. On average, transcription precedes peak transcript abundance by approximately 80 minutes [3]. Although transcript stabilization has been reported to increase during the schizont stage [29], this phenomenon appears to affect only a relatively small subset of genes, whereas the majority of genes undergo RNA decay at comparable rates throughout the IDC [3]. Consequently, the temporal patterns described here likely provide a close approximation of the underlying transcriptional program. Overall, our findings are consistent with the ‘just-in-time’ model of gene expression, in which genes are transcribed shortly before their products are required for stage-specific biological processes.

The gene expression profiles observed during schizogony closely mirror the timeline of merozoite organelle biogenesis established by ultrastructural and fluorescence imaging studies [30–32]. Of the three classes of secretory organelles present in merozoites, rhoptries form first in mid-stage schizonts, consistent with our finding that rhoptry-associated genes reach peak expression between 42 and 45 hpi. Micronemes are assembled next, coinciding with the onset of merozoites segmentation [32]. Distinct subsets of micronemes have been described, with AMA1-containing micronemes positioned more apically than those containing EBA-175 [31–34].

In line with this spatial organization, AMA1 reaches peak expression earlier than EBA-175 (**Table S2**), suggesting that temporal differences in gene expression are resolved even among genes encoding proteins destined for the same type of organelle. Dense granules are formed last and are fully established in mature segmented merozoites [32]. Dense granules secrete their contents into the nascent parasitophorous vacuole after the parasite has entered the erythrocyte [35], and contain PV1 and components of the PTEX translocon [36–38], in agreement with the late expression of dense granule-associated genes observed in our data set. Together, these findings suggest that the transcriptional program is coordinated with merozoite organelle biogenesis, potentially contributing to the orderly sorting and trafficking of proteins to their appropriate secretory organelles, in addition to established mechanisms such as targeting motifs and molecular chaperones [39,40].

Our analysis of transcription factors associated with invasion gene expression suggests that these gene modules are regulated through the cooperative action of multiple ApiAP2 transcription factors rather than by a single master regulator. Among the different modules, microneme genes were notable for their apparent lack of targeting by AP2-I, AP2-LT, and AP2-P. However, our analysis was limited to the four ApiAP2 transcription factors for which ChIP-seq data are currently available in *P. falciparum*. In *P. berghei*, it has been shown that AP2-O is involved in regulation of microneme gene expression in ookinetes and that deletion of AP2-O attenuates growth of blood-stage parasites [25,26]. Furthermore, a recent study identified ApiAP2 transcription factor SIP2 as a regulator of invasion gene expression in *P. berghei* [41]. Because the mapping of invasion gene orthologs between *P. falciparum* and *P. berghei* is incomplete, we were unable to assess the contribution of SIP2 and AP2-O to the regulation of the distinct invasion gene modules. The transcriptional regulatory network governing invasion gene expression is therefore likely to be more extensive and complex than our analysis suggests. A further limitation is that the available ChIP-seq datasets were not optimally timed for resolving regulatory interactions during the final stages of schizogony. Most experiments were performed between 30 and 40 hpi, well before the peak expression of many late schizont genes. Consequently, binding events occurring immediately prior to egress may have been missed, potentially explaining the apparent absence of AP2-I, AP2-LT, and AP2-P binding at microneme-associated genes, although the same limitation applies to the AP2-EXP data set. Future studies will be needed to further resolve the transcriptional regulatory network controlling invasion gene expression by determining the contributions of SIP2 and AP2-O and elucidating how multiple ApiAP2 transcription factors cooperate to regulate distinct invasion gene modules.

In addition to delineating the transcriptomic landscape of merozoite formation, our findings provide a more precise timing of gene expression during the schizont-to-ring transition, enabling more accurate assignment of parasite genes to pre-egress and post-invasion developmental programs. As one example, we found that transcripts encoding ETRAMP2, ETRAMP11.1, ETRAMP11.2, and ETRAMP12 reach peak abundance in late schizonts, whereas the closely related genes encoding ETRAMP10.1 and ETRAMP14.1 are maximally expressed in ring-stage parasites. This sequential pattern is in agreement with the findings of Spielmann *et al.*, who detected expression of ETRAMP2, 11.1, 11.2, and 12 in parasites harvested at 40 – 4 hpi, while ETRAMP10.1 and ETRAMP14.1 transcripts first appeared during the early ring stage (0 – 6 hpi) [42]. This example illustrates that the temporal resolution afforded by our scRNA-seq atlas enables the discrimination of distinct expression programs among closely related gene family members, which provides direction for investigating their functional specialization and the regulatory mechanisms that coordinate their expression.

In our prior work, we analyzed the transcriptional landscape in merozoites isolated following E64 treatment [12]. The use of an egress inhibitor was necessary to isolate sufficient numbers of merozoites, as the extracellular phase of the IDC is very short. Under *in vitro* conditions, the interval between egress and invasion averages approximately 1.5 minutes [43]. The invasion potential of free merozoites decreases within minutes following isolation [11], suggesting that merozoites are fragile and do not remain invasive for prolonged period of time. In line with the kinetics of the invasion process, we found that free merozoites rapidly show signs of cellular damage, including relatively high mitochondrial mRNA and low cytoplasmic mRNA content. Our results showing deterioration of E64-treated schizonts and merozoites are seemingly at odds with the isolation of viable merozoites through mechanical rupture of E64-treated schizonts [10,11]. However, invasion efficiencies under such conditions generally do not exceed 1 – 2%, although invasion rates of 15 – 18% can be reached under optimal conditions [11]. The large majority of merozoites obtained following E64 treatment thus fail to invade and may disintegrate rapidly, consistent with the transcriptional profile we describe here.

Our results demonstrate that PIPseq is a suitable platform for single-cell transcriptomic profiling of *P. falciparum* blood-stage parasites. The mixed-stage PIPseq dataset generated in this study showed improved representation of late-stage schizonts as compared with a publicly available mixed-stage 10x Genomics dataset. However, these datasets were generated in different laboratories from independent parasite cultures and therefore do not permit a controlled head-to-head comparison of platform performance. In addition, both libraries were produced using earlier generations of their respective chemistries. Because single-cell RNA-seq chemistries continue to evolve rapidly, direct comparisons between platforms can quickly become outdated as technologies improve. Consequently, our findings should be interpreted as evidence that PIPseq can effectively capture the full spectrum of *P. falciparum* blood-stage development, including the sparsely represented late schizont stages, rather than as a formal benchmark against 10x Genomics. However, advantages of the PIPseq workflow over the 10x Genomics platform include its affordability, scalability, and the ease of sample preparation that allows for sampling across time courses to achieve fine-scale temporal resolution.

In summary, our study provides a high-resolution scRNA-seq atlas of the *P. falciparum* schizont-to-ring transition, capturing the coordinated transcriptional programs that underlie merozoite maturation, egress, invasion, and early intracellular development. By resolving the temporal ordering of gene expression during this developmental window, we define coordinated transcriptional programs required for organelle biogenesis, invasion, and establishment of the intracellular niche. In addition, we identify transcriptional changes associated with loss of merozoite viability during prolonged extracellular residence. Our data provides a resource for deeper analysis of parasite gene function and regulatory networks during one of the most dynamic and experimentally inaccessible stages of the IDC.

## MATERIALS & METHODS

### Ethics approval statement

Blood for *P. falciparum* cultures was obtained from local blood donors recruited at the University of Texas Health Science Center at San Antonio. Donors provided verbal informed consent for the use of their blood for research. This protocol was approved by the Institutional Review Board of the University of Texas Health Science Center at San Antonio (protocol 20230283HU).

### Parasite culture

*P. falciparum* strain 3D7 parasites were cultured [44] in human O^+^ erythrocytes at 3 – 10% parasitemia in complete culture medium (5% hematocrit). Complete culture medium consisted of RPMI 1640 medium (Gibco #32404014) supplemented with gentamicin (45 µg/mL final concentration; Gibco #15710064), HEPES (40 mM; Fisher #BP3101), NaHCO_3_ (1.9 mg/mL; Sigma #SX03201), NaOH (2.7 mM; Fisher #SS266-1), hypoxanthine (17 µg/mL; Alfa Aesar #A11481-06), L-glutamine (2.1 mM; Corning #25005Cl), D-glucose (2.1 mg/mL; Fisher #D16-1), and 10% heat-inactivated human AB^+^ serum (Valley Biomedical #HP1022). Parasites were cultured at 37°C in an atmosphere of 5% O_2_, 5% CO_2_, and 90% N_2_ under continuous agitation [45] (19 mm orbital diameter; 50 rpm for 25 cm^2^ flasks; 40 rpm for 75 cm^2^ flasks). Before use in cultures, 12.5 mL of freshly isolated packed erythrocytes were washed twice with 10 mL cold incomplete medium (complete culture medium without human serum) and pelleted between each wash by centrifugation at 500 × *g* for 8 min. at 4°C (max. acceleration and slow break; Beckman Coulter Allegra X-14R). Washed erythrocytes were resuspended in 1 volume of complete medium to 50% hematocrit, stored at 4°C, and used within two weeks of collection.

Parasites were synchronized to the ring stage by treatment with 5% D-sorbitol (Fisher #S459-500) [46]. Cultures containing 5 – 10% of ring-stage parasites were centrifuged at 250 × *g* for 5 min. at RT. Pelleted erythrocytes were resuspended in 10 volumes of 5% D-sorbitol in MQ water, vortexed for 30 sec. and incubated for 8 min. at 37°C. The cells were vortexed another 15 sec., washed with 5 volumes of complete culture medium (250 × *g* for 5 min. at RT), resuspended in complete culture medium at 5% hematocrit, and cultured as described above.

### Preparation of PIPseq libraries

Parasites were processed for single-cell sequencing using the PIPseq T20 3’ Single Cell RNA Kit v3.0 (Fluent Biosciences #FBS-SCR-T20-4-V3) with a targeted recovery of 20,000 cells and <8% multiplet rate or the PIPseq T2 3’ Single Cell RNA Kit v4.0PLUS (Fluent Biosciences #FBS-SCR-T2-8-V4.05) with a targeted recovery of 2,000 cells and ∼5% multiplet rate, as specified for individual libraries below. Uninfected erythrocytes in the samples were not counted towards the number of cells processed for capture and library preparation. All libraries were sequenced on a NovaSeq X Plus (Illumina) in a 150PE run.

For mixed-stage parasites, 40 µL of culture at 5% parasitemia was added to 1 mL of pre-warmed (37°C) Cell Suspension Buffer (CSB, provided in the kit), centrifuged at 250 × *g* for 5 min. at RT, and resuspended in 400 µL cold (4°C) CSB. Of this suspension, 20 µL (corresponding to approximately 40,000 infected erythrocytes) was immediately added to the PIP reaction. This sample was further processed using the T20 kit according to the manufacturer’s instructions.

To generate the library enriched for schizonts, merozoites, and early ring-stage parasites (SZ-MZ-ER), a culture of *P. falciparum* 3D7 parasites was treated with sorbitol to remove all trophozoites and schizonts. After 42 hours, 16 mL of culture (4% ring-stage and 6% mature parasites) was applied to two LS columns (Miltenyi #130-042-401) attached to a Midi-MACS separator (Miltenyi #130–042-501) to isolate mature parasites. In brief, each column was first equilibrated with 5 mL complete medium. The culture was centrifuged at 250 × g for 5 min. at RT to pellet erythrocytes. The pellet was resuspended in complete medium to 30% hematocrit and split over the two columns. The flowthrough was collected and then passed back through the columns to further capture mature parasite-infected erythrocytes. Each column was washed with 2.75 mL complete medium before taking it off the magnet and applying 1 mL complete medium to elute the bound infected erythrocytes. The isolation was repeated with the flowthrough and the first 0.75 mL of the wash. Together, these two treatments resulted in a synchronized batch of parasites that were approximately between 42 and 48 hours post-invasion (hpi). All elutions (4 mL total) were combined, before splitting it into two separate cultures. The first culture (2 mL) was maintained without adding uninfected erythrocytes for the purpose of isolating free merozoites. To the second culture (2 mL), 200 μL uninfected erythrocytes at 50% hematocrit were added to allow invasion and ring-stage development to occur. Both cultures were returned to the incubator for four hours. Next, the first culture was used for the isolation of naturally egressed merozoites. The culture was centrifuged at 250 × *g* for 5 min. at RT to pellet late schizonts and remove any existing free merozoites in the culture supernatant. The schizonts were resuspended in culture medium and incubated at 37°C for 10 min. to allow for merozoite egress. Next, the unruptured schizonts were pelleted again at 250 × *g* for 5 min. at RT, while free merozoites remained in the supernatant. The supernatant was then transferred to a new tube and centrifuged at 4,000 × g for 5 min. at RT to pellet the merozoites. At this point, most of the parasites in the second culture were expected to be a mix of late schizonts (between 46 and 48 hpi) and early ring-stage parasites (between 0 and 4 hpi). Of this culture, at approximately 3% parasitemia, 40 µL was added to 1 mL pre-warmed CSB, followed by centrifugation (250 × *g* for 5 min. at RT). The cell pellet was resuspended in 800 µL cold CSB and 20 µL of this suspension (approximately 12,000 infected erythrocytes) was added to the pelleted merozoites. This cell suspension (20 µL), containing schizonts, merozoites, and rings, was added to the PIP reaction and processed for PIPseq using the T20 kit according to the manufacturer’s instructions.

For the E64 SZ and E64 MZ samples, parasite cultures underwent double sorbitol treatment, 14 hours apart, to obtain tightly synchronized parasites. The following day, infected erythrocytes containing parasites in the late-trophozoite and schizont stages were isolated from culture by magnetic separation as described above. Infected erythrocytes were eluted from the column in a total of 3 mL complete culture medium. Infected erythrocytes were then incubated with E64 (10 µM final concentration, Sigma #324890-1MG) for 6 hours at culture conditions described above to allow the parasites to develop into fully segmented schizonts while preventing egress from the erythrocytes. A second culture containing 2.7 mL eluted infected erythrocytes and 0.3 mL uninfected erythrocytes at 50% hematocrit was maintained in parallel. Parasitemia of this flask was determined by thin smear to estimate the concentration of infected erythrocytes in the E64 treatment flask. E64-treated schizonts (40 µL) were added to 1 mL pre-warmed CSB, centrifuged (200 × *g* for 3 min. at RT), and resuspended in 640 µL cold CSB. An estimated 5,000 infected erythrocytes (5 µL) were added to the PIP reaction and processed using the T2 kit according to the manufacturer’s instructions. To obtain E64-treated merozoites, the remaining culture was pelleted by centrifugation at 1,900 × *g* for 8 min. at RT and the supernatant containing E64 was removed. The pellet was resuspended in 2 mL warm (37°C) incomplete medium. Merozoites were released from the erythrocytes by passing them through a 1.2 µm syringe filter (Pall #4190) and were subsequently run over a magnetic column to remove free hemozoin and any residual schizonts. Merozoites were then pelleted by centrifugation at 4,000 × *g* for 5 min. at RT, washed in pre-warmed CSB, and finally resuspended in cold CSB. An estimated 5,000 merozoites (5 µL) were added to the PIPs and processed using the T2 kit according to the manufacturer’s instructions.

### Single-cell RNA-seq data analysis

Sequencing reads were mapped to the *P. falciparum* strain 3D7 reference genome (PlasmoDB v64) [47,48] and assigned to individual cells of origin using PIPSeeker (Fluent Biosciences) version 2.1. Results filtered at the highest sensitivity (sensitivity 5) were then processed using Seurat version 5.3.1 [49] in RStudio (v2025.09.2+418) with R (v4.5.1), following the default Seurat v4 pipeline. The 10x Genomics *P. falciparum* data set of asexual stages was downloaded from the Malaria Cell Atlas (*P. falciparum* 10x set1) [14]. Low-quality cells with many (≥1,000; indicative of multiplets) or few (≤100; indicative of dying cells or empty droplets that captured ambient RNA) genes detected were filtered from the samples. For the SZ-MZ-ER library, we were unable to estimate the number of merozoites added to the sample prior to library preparation due to time constraints. As a result, the total number of cells recovered from this library was >50,000 and included many multiplets. To prevent this sample from dominating the analysis, the data set was downsampled to 5,000 cells.

Raw unique molecular identifier (UMI) counts were normalized by the total UMI count of each cell, multiplied by a scale factor (10,000), and log-transformed using Seurat’s *LogNormalize* method. For each library, we determined the 2,000 most variable genes among all cells within that sample based on their average expression and dispersion using the default method (variance-stabilizing transformation), followed by scaling and centering all features in the data set. Principal component analysis was performed on the 2,000 most variable genes to reduce the dimensionality of the data. When integrating multiple data sets, integration anchors were identified using the function *FindIntegrationAnchors*, and data sets were combined using the *IntegrateData* function. Scaling and principle component analysis were then performed again on this combined object.

To determine the number of dimensions to use for visualization and clustering, we calculated two potential cutoffs. The first cutoff was the first principal component (PC) where the cumulative explained variance exceeded 90% and the PC individually explained less than 5% of variance. The second cutoff was the first PC where the difference in explained variance between consecutive PCs was less than 0.1 percentage points. The smallest of these two cutoff values was subsequently used as the number of dimensions in the *RunUMAP* and *FindNeighbors* functions. We then performed clustering using the Louvain algorithm implemented in the *FindClusters* function of Seurat (algorithm = 1; resolution = 0.5 for UMAP #1 and #2, resolution = 0.9 for UMAP #3 intermediate, resolution = 0.3 for the final UMAP #3, and resolution = 0.7 for the high-resolution version of the final UMAP #3). Because cells from all samples were distributed evenly across the UMAPs, no batch corrections were performed. For integrated data sets, the default assay was switched back to ‘RNA’ following cell clustering, and the layers of the various data sets were joined using the *JoinLayers* function.

Clusters corresponding to gametocytes, multiplets (in particular in the SZ-MZ-ER library), and other small outlier clusters were removed and the remaining clusters were renumbered to represent the developmental progression through the intraerythrocytic developmental cycle. UMAP plots with cluster assignments or overlaid with gene expression levels, as well as violin plots of gene expression, were generated using standard Seurat functions. **Table S7** contains a list of gene IDs of individual genes plotted in the main and supplementary figures. All R code used to analyze and visualize scRNA-seq data is provided in **Files S1 - 4**.

### Transcriptome complexity analysis

To analyze the number of transcripts and the number of genes detected per cell, we extracted the metadata from the Seurat object. The two relevant metrics, nCount_RNA and nFeature_RNA, were then plotted by cluster using the R package ggplot2 [50]. To assess whether the variation in detected genes in cells from different clusters showed different patterns even after accounting for sequencing depth, nFeature_RNA was modeled as a function of nCount_RNA and cluster membership in the Seurat metadata. Because the relationship between these metrics was nonlinear on the raw scale, both variables were transformed using log1p prior to analysis. We first fit a depth-only linear regression model (log1p(nFeature_RNA) ∼ log1p(nCount_RNA)) and then fit a second model that additionally included cluster identity as a categorical predictor (log1p(nFeature_RNA) ∼ log1p(nCount_RNA) + cluster). Cluster-specific residual differences from both models were plotted to determine to what extent the models explained variance in detected gene counts.

### Analysis of peak gene expression

The aggregated counts per cluster were extracted from the Seurat object using the *AggregateExpression* function and stored in a new object. These counts were normalized using the *NormalizeData* function and subsequently extracted using the *GetAssayData* function. Log-normalized gene expression values for all genes in all clusters in UMAPs #1, #2, and #3 (intermediate, final, and high-resolution) are provided in **Table S2**. To remove genes with low expression levels, only genes with a minimum log-normalized expression value of 0.2 in any cluster were retained. For each gene, peak expression was defined as the cluster with the highest log-normalized gene expression value. Since UMAP #3 contained only a subset of cells, this analysis was restricted to genes with peak expression in the relevant developmental stages by including only genes that peaked in expression in their original clusters (0, 2, 4, 12, and 17) in the intermediate UMAP #3 containing all cells (**Fig. S9**). For visualization purposes, heatmaps plotting gene expression values were sorted based on the cluster of peak expression and generated using the pheatmap package in R [51].

### Differential gene expression analysis

To identify differentially expressed genes between the normal developmental trajectory and the diverging branch in UMAP #3, we used the *FindMarkers* function from Seurat with ident.1 set to clusters 5 – 7 and ident.2 set to clusters 1 – 4, and logfc.threshold = 1. Cluster-specific marker genes in UMAP #3 HR were identified using the *FindAllMarkers* function with parameters only.pos = TRUE, min.pct = 0.25, logfc.threshold = 0.25. Both functions were performed using the default Wilcoxon rank sum test with Bonferroni correction for multiple testing. Results were filtered for adjusted P value < 0.05 and only genes with a minimum log-normalized expression value of 0.2 in any cluster of the relevant data set were retained.

### Gene set analysis

The expression of gene modules was calculated by averaging the z-scored log-normalized expression of genes within each gene set per cluster. For the analysis of invasion gene sets previously defined by Poran *et al*. [17], a new module of merozoite surface proteins was added by filtering *P. falciparum* gene descriptions on “merozoite surface protein”. For the gene sets expressed during the schizont-to-ring transition, the sums of unique peptides were extracted from two independent proteomics studies of merozoites (ref. [38] and Shahid Kahn, unpublished, downloaded from PlasmoDB).

### ApiAP2 transcription factor target gene analysis

Lists of target genes of ApiAP2 transcription factors AP2-I, AP2-LT, AP2-P, and AP2-EXP were retrieved from publications reporting the results of ChIP-seq experiments performed using *P. falciparum* asexual blood-stage parasites at 30 – 40 hpi [21–24]. Target genes were matched with genes in each of the schizont-stage gene modules (**Table S3**) and the UMAP #2 clusters (**Table S2**) to perform the analyses described in the text.

### DNA motif search

To search for enrichment of DNA motifs in gene promoter regions, the coordinates of the regions 500 nucleotides upstream of the gene start sites were extracted from the *P. falciparum* strain 3D7 gene annotation file (PlasmoDB v68) and stored in bed format. DNA sequences corresponding to these regions were extracted from the *P. falciparum* strain 3D7 genome using the *homerTools* function in the Homer package [52]. The homer function *findMotifsGenome.pl* was used to identify DNA motifs of 6 or 8 nucleotides long.

### Gene ontology enrichment analysis

All nuclear genes with higher expression in clusters 5 – 7 relative to clusters 1 – 4 of UMAP #3 were extracted (n = 342). Gene ontology enrichment analysis was performed using PANTHER through the Gene Ontology knowledgebase [53–55]. Terms that were significantly enriched (Fisher’s exact test, False Discovery Rate < 0.05) are presented in **Table S5**.

## ACKNOWLEDGEMENTS

This work was supported by a pilot project award from the Department of Microbiology, Immunology, and Molecular Genetics at the University of Texas Health Science Center at San Antonio. The authors acknowledge the Texas Advanced Computing Center (TACC) at The University of Texas at Austin for providing computational resources that have contributed to the research results reported within this paper. The following reagent was obtained through BEI Resources, NIAID, NIH: *Plasmodium falciparum*, Strain 3D7, MRA-102, contributed by Dr. Daniel J. Carucci.

## AUTHOR CONTRIBUTIONS

Conceptualization: Ian H. Cheeseman, Evelien M. Bunnik

Formal analysis: Jake A. Moore, Evelien M. Bunnik

Funding acquisition: Ian H. Cheeseman, Evelien M. Bunnik

Investigation: Sebastiaan Bol, Jake A. Moore, Evelien M. Bunnik

Project administration: Evelien M. Bunnik

Supervision: Evelien M. Bunnik

Visualization: Evelien M. Bunnik

Writing – original draft: Sebastiaan Bol, Siddharth Neog, Evelien M. Bunnik

Writing – review & editing: Sebastiaan Bol, Siddharth Neog, Vinit Upasani, Ian H. Cheeseman, Evelien M. Bunnik

## DATA AVAILABILITY

Raw and processed single-cell sequencing files are available from the NCBI Gene Expression Omnibus under accession no. GSE339432. All other relevant data and the custom code used for data analyses are within the paper and its supporting information files. Correspondence should be addressed to Evelien M. Bunnik.

## COPYRIGHT AND PERMISSION STATEMENT

No copyrighted, trademarked, and reproduced material is included in this article.

